# High-fat diet suppresses the positive effect of creatine supplementation on skeletal muscle function by reducing protein expression of IGF-PI3K-AKT-mTOR pathway

**DOI:** 10.1101/347120

**Authors:** Renato Ferretti, Eliezer Guimarães Moura, Veridiana Carvalho dos Santos, Eduardo José Caldeira, Marcelo Conte, Cintia Yuri Matsumura, Adriana Pertille, Matias Mosqueira

**Author notes:** Corresponding Authors (RF), (MM). These authors contributed equally to this work (Renato Ferretti, Matias Mosqueira). These authors also contributed equally to this work (Eliezer Guimarães Moura, Veridiana Carvalho dos Santos).

## Abstract

High-fat (HF) diets in combination with sedentary lifestyle represent one of the major public health concerns predisposing to obesity and diabetes leading to skeletal muscle atrophy, decreased fiber diameter and muscle mass with accumulation of fat tissue resulting in loss of muscle strength. One strategy to overcome the maleficent effects of HF diet is resistance training, a strategy used to improve muscle mass, reverting the negative effects on obesity-related changes in skeletal muscle. Together with resistance training, supplementation with creatine monohydrate (CrM) in the diet has been used to improve muscle mass and strength. Creatine is a non-essential amino acid that is directly involved in the cross-bridge cycle providing a phosphate group to ADP during the initiation of muscle contraction. Besides its antioxidant and anti-inflammatory effects CrM also upregulates IGF-1 resulting in hyperthophy with an increase in muscle function. However, it is unknown whether CrM supplementation during resistance training would revert the negative effects of high-fat diet on the muscle performance. During 8 weeks we measured muscle performance to climb a 1.1m and 80° ladder with increasing load on trained rats that had received standard diet or high-fat diet, supplemented or not with CrM. We observed that the CrM supplementation up-regulated IGF-1 and phospho-AKT protein levels, suggesting an activation of the IGF1-PI3K-Akt/PKB-mTOR pathway. Moreover, despite the CrM supplementation, HF diet down-regulated several proteins of the IGF1-PI3K-Akt/PKB-mTOR pathway, suggesting that diet lipid content is crucial to maintain or improve muscle function during resistance training.

## Introduction

High-fat (HF) diets and sedentary lifestyle represent a public health concern that can predispose to obesity and diabetes [1], which can lead to skeletal muscle atrophy due to degradation of muscle fibers, a reduction of fiber type 1 (aerobic metabolism) and with an increase of the fiber type 2X (glycolic metabolism) [2, 3]. Obesity is further characterized by the loss of muscle strength, decreased fiber diameter and muscle mass with accumulation of fat tissue [4]. The skeletal muscle constitutes about 40- 50% of body mass and is the main responsive tissue to insulin-stimulated uptake of glucose and fatty acids. At the cellular level, HF diets induce mitochondrial dysfunction leading to insulin resistance and reducing the muscle mass via decreasing protein levels of the IGF1-PI3K-Akt/PKB-mTOR skeletal muscle growth pathway, i.e. the insulin receptor substrate 1 (IRS1), phosphoinositide 3-kinase (PI3K), and a serine-threonine protein kinase (AKT) [5]. Moreover, obesity upregulates myostatin (GDF-8), a member of the transforming growth factor-β (TGF-β1) family, FoxO, inducible nitric oxide synthase and Csp3; all members of the muscle atrophy pathway [3, 6]. Although, both anabolic and catabolic pathways are well described, i.e., AKT inhibits FoxO and myostatin-SMAD 2/3 inhibits AKT [3], the exact regulation of protein metabolism during obesity is still incompletely characterized [7].

Resistance training as one strategic treatment of physical rehabilitation against obesity results in increased force-generation capacity, improved muscle mass and positive effects on obesity-related changes in skeletal muscle [8]. This technique increases the expression of IGF-1, which in the mouse decreases myosin 2B expression and increases myosin 2X expression, while in humans, there is a downregulation of the fast 2X myosin and an upregulation of myosin 2A [2].

One method to enhance the effectiveness of resistance training is supplementation with creatine monohydrate (CrM) in the diet. CrM has been used not only for athletes as an ergogenic aid for improving muscle mass and strength, but also as therapeutic agent for patients suffering from sarcopenia, muscle wasting and myopathies [9-11]. Creatine is a non-essential amino acid that is synthetized in the liver and kidney or ingested from the meat or artificial supplements. In the muscle, creatine is found as free creatine or phosphocreatine, both are directly involved in cross-bridge cycling providing phosphate groups to ADP during initiation of muscle contraction [11]. CrM has an antioxidant and an anti-inflammatory effect, reducing lipid peroxidation and DNA susceptibility to oxidative stress [12–15]. It has also been shown that CrM upregulates IGF-1 in cultured myotubes [16] and in human skeletal muscle resulting in hypertrophy with increased muscle function [10, 17, 18]. The precise mechanism by which CrM upregulates IGF1 and thus the differentiation of myogenic muscle fibers and hypertrophy remains unknown. The strategy to use CrM supplementation in obesity associated with hypertrophic response during resistance training has produced positive results depending on the dosage, duration of the treatment and on the type of physical training [10]. Therefore, there are still lacunae regarding the effect of CrM supplement on the muscle performance during resistance training during high-fat diet. To test this hypothesis, we compared the muscle capacity of trained rats who received standard or high-fat diet, supplemented with CrM or not to climb a ladder (1.1m high at 80° incline) with increasing loads during 8 weeks. We observed that the improvement of muscle performance seen in trained rats receiving standard diet supplemented with CrM was completely canceled under the HF diet.

## Materials and methods

### Care and use of animals

Forty male Wistar rats (HanUnib; *Rattus novergucis*) were obtained from the breeding colony at the State University of Campinas (CEMIB-UNICAMP) and maintained by our institutional animal care facility. The rats were kept in collective cages (2- 3 animals per cage) at constant temperature (21 ± 2°C), cycles of 12h light/ 12h darkness and with free access to food and water. All animal procedures were performed in accordance with the Guide for Care and Use of Laboratory Animals. The committee of experimental animal approved the protocol CEUA#490/2012.

### Experimental groups

At the time of the experiments, all animals were 24 weeks of age and were randomized into the following eight experimental groups according to their diet, training and creatine supplementation: i) untrained (UT) standard diet (SD), ii) untrained creatine supplemented (SD-CrM), iii) resistance training (SD-T), iv) resistance training with creatine supplementation (SD-T-CrM), v) untrained high-fat diet (HF), vi) untrained HF with creatine supplemented (HF-CrM), vii) HF and resistance training (HF-T) and viii) HF and resistance training with creatine supplementation (HF-T-CrM).

### Diet

The animals from the SD, SD-CrM, SD-T and SD-T-CrM received standard diet (Nuvital, Nuvilab, Brazil) containing 71g of carbohydrate, 23g of protein, 6g of total fat and 5g of fiber, totaling 3.8 kcal/g. Eight weeks prior to the beginning of the experiments and during the eight weeks of experimental procedures, obesity rats groups (HF, HF-CrM, HF-T and HF-T-CrM) received a high-fat diet (Nuvital, Nuvilab, Brazil) containing 38g of carbohydrate, 15g of protein, 46g of total fat and 5g of fiber, totalling 5.4 kcal/g. Animals had free access to water and chow during the experimental period. The CrM supplementation was given daily from day 1 until the last day of the experimental procedure.

### Resistance-training protocol

At week 15 prior to the beginning of the experiments, the resistance training groups (SD-T, SD-T-CrM, HF-T and HF-T-CrM) were submitted to climbing sessions three times per week during 8 weeks, according to Cassilhas and co-workers[19]. The rats were adapted to climbing a vertical ladder (1.1 × 0.18m, 2cm grid, 80° of inclination) with weight attached to falcon cylinder clipped to the base of the tail wrapped with paper hypoallergenic tape (3M™Micropore™). The length of the ladder lead to 8-12 movements per climb. A three-day adaptation was performed one week before the training session. The climbing training consisted of two introductory climbs, followed by three full length climbing attempts. First, the animal was placed at the top of the ladder near the resting area (40×20×20 cm). Rats were motivated to climb by a touch to the tail with tweezers. Second, rats were positioned in the middle of the ladder and an external stimulus was applied to encourage climbing. Finally, during the following three full length climbing attempts, rats climbed from the base of the equipment to the ladder’s top. The rats that refused to climb were excluded. The adaptation climbing was done using only the body weight. The first training session started two days after the adaptation period with 50% of the body weight attached to each animal. A series of 30g weights were added until the maximal load encumbered the rat’s capacity to climb and consisted of four to twelve ladder climbs. After every successful climbing from the bottom to the ladder’s top, the rats were allowed to rest for 120 seconds. Failure was defined after three non-successful attempts. The maximal carrying load was considered the highest load before the failed attempts. The training session consisted of four climbs with 50%, 75%, 90% and 100% of the rat’s maximal carrying load. After each fourth climb, additional 30g weights were added until the new maximal carrying load was determined.

### Quantitative analysis of training

Maximal carrying load was determined by the total amount of load carried to the top of the ladder. The total isotonic contraction measured in grams was calculated by summing the body weight and the weight lifted to the top of the ladder times the number of repetitions (number of times the rat successfully climbed to the top of the ladder). Work measured in kilo Joule was calculated multiplying total mass lifted to the top of the ladder, the length of the ladder (1.1m), gravitational force (9.8 06 ms^−2^) and the ladder’s angle (sen80 = 0.9848).

### Sample collection and tissue preparation

After the rats rested for 48h after the last climbing session, they were anesthetized with a mix of ketamine (80 mg/kg of body weight) and xylazine (12 mg/kg of body weight) and left and right gastrocnemius were rapidly dissected and one was snap frozen in N-hexane cooled in liquid nitrogen, and stored at −80°C. The frozen muscles were transversal cross-sectioned (8-μm thick cryostat sections), and then stained with heamtoxylin-eosin (HE) for histological analysis, where the cross sectional area and Feret’s fiber diameter was calculated using Image J 1.51f software (National Institute of Health, USA). Fiber sizes from each experimental condition were determined from 5-7 randomly captured images.

### Determination of protein levels

Muscle samples from the second gastrocnemius of each rat were lysed in assay lysis buffer containing freshly added protease and phosphatase inhibitors (1% Triton X-100, 100 mM Tris-HCl, pH 7.4, 100 mM sodium pyrophosphate, 100 mM NaF, 10 mM sodium ortho-vanadium, 10 mM EDTA, 2 mM PMSF, and 10 μg/ml aprotinin). The samples were centrifuged for 20 min at 11,000 rpm, and the soluble fraction was resuspended in 50 μl Laemmli loading buffer (2% SDS, 20% glycerol, 0.04 mg/ml bromophenol blue, 0.12 M Tris-HCl, pH 6.8, and 0.28 M β-mercaptoethanol). Samples were stored at −80°C until the analysis. The proteins were resolved on 8%– 12% SDS-polyacrylamide gels and transferred to a nitrocellulose membrane. Primary antibodies were diluted in TBS containing 0.05% Tween (TBS-T). Membranes were incubated overnight with primary antibodies at 4 °C (S1-Table 1). For secondary antibody incubation, anti-rabbit or anti-mouse HRP (Promega) were diluted in TBS-T containing 5% skim milk (S1-Table 1). Results were visualized with enhanced chemiluminescence (ECL) SuperSignal West Pico Chemiluminescent Substrate kit (Pierce Biotechnology). For protein loading control, the blots were stripped and re-probed for glyceraldehyde-3-phosphate dehydrogenase (GAPDH). Band intensities were quantified using ImageJ 1.38X (National Institute of Health, USA) software.

### Statistical analysis

All statistical analyses were performed using GraphPad Prism version 6 for Windows, GraphPad Software, La Jolla California USA, www.graphpad.com. One-way analysis for variance (ANOVA) was used with a *post hoc* multiple-comparison. Sidak’s multiple comparison test was used on body weight, epididymal fat mass, lean body mass and protein levels. Multiple unpaired two-tailed Student’s t test for pairwise comparison utilizing Benjamin & Hochberg’s method and the FDR (Q) = 5% was used on Maximal carrying load, total isotonic contraction, work and relative carrying load. Non-parametric one-way ANOVA Kruskal–Wallis test was used on CSA and Minimal Feret’s diameter. Significance was considered as p < 0.05.

## Results

### HF diet increases body weight and epididymal fat mass

The effect of high-fat (HF) diet was evaluated measuring body weight (Fig 1A) and epididymal fat mass (Fig 1B). Two-way ANOVA showed that there was no interaction between exercise and diet and no differences within the exercise parameters; however, the diet parameter was highly significantly different (F_1,31_ = 82.04; p< 0.001). Based on these results, we pairwise compared the effect of diet on the exercise treatment, observing that in comparison to the standard diet (SD), HF increased significantly body weight in all four treatments: untrained (UT), trained (T), diet supplemented with creatine monohydrate (CrM) and the combination of T and CrM (T-CrM). Next, we evaluated whether this effect was due to an increase in epididymal fat mass or just a proportional increase of the body (Fig 1A, S1-Table 2). The two-way ANOVA showed that again diet was the only significant parameter (F_1,33_) = 186.5; p< 0.001. Once more, the post-hoc pairwise comparison between SD and HF diets showed that the increase of the epididymal fat mass was independent of the exercise treatment (Fig 1B, S1-Table 3).

**Figure 1.**
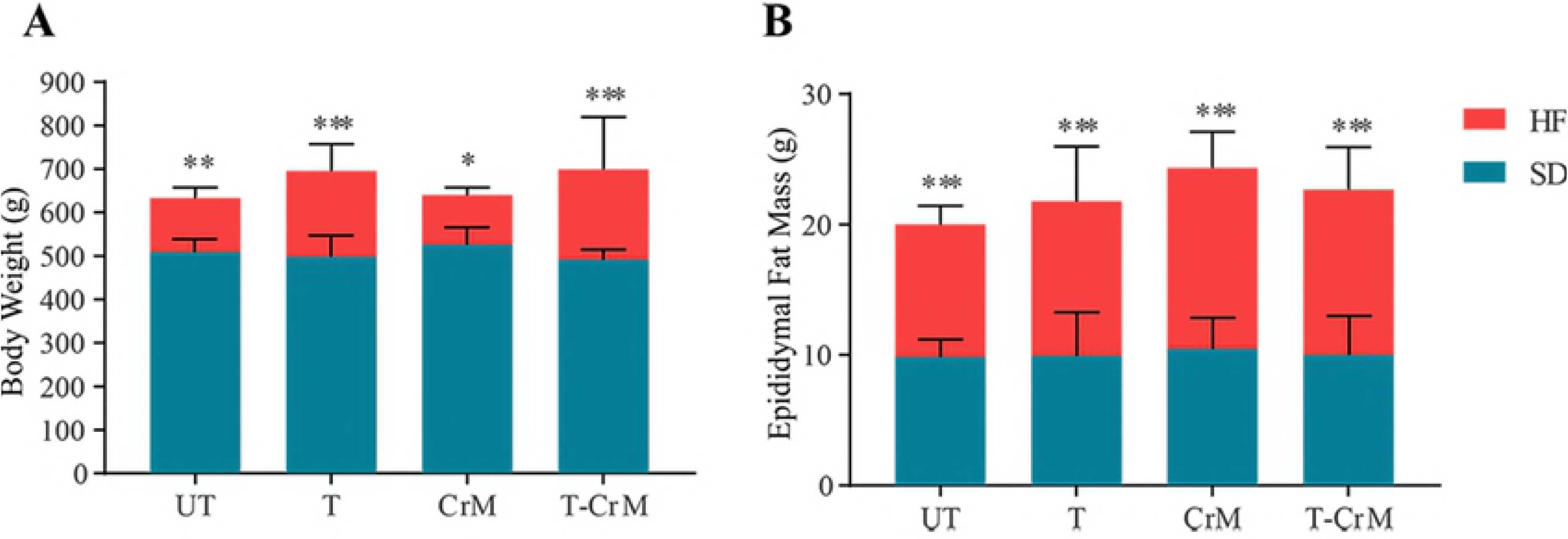
Effect of high-fat diet on the body mass of rats. A. Comparison between standard diet (SD, (blue) and the administration of a high-fat diet (HF; red) on rat’s body weight (g) in untrained (UT) rats, trained (T) rats, with creatine monohydrate supplementation (CrM) and in trained rats with CrM supplementation (T-CrM). B. The effect of HF diet on the rat’s epididymal fat mass (g) in comparison to SD after UT, T, CrM and T-CrM treatments. Response of the rat’s lean body mass (g) on HF diet in comparison to SD under UT, T, CrM and T-CrM treatments. n= five rats in each group; * p< 0.05; ** p< 0.01; *** p< 0.001.

### CrM supplementation improves muscle performance in trained rats

Since diet has the major effect on the previous evaluated parameters and knowing the benefits of CrM supplementation of muscle performance, we evaluated whether CrM supplementation is able to change muscle performance overriding the dietary effect on trained rats. In order to measure the dietary effect on muscle performance over time and reducing the number of animals necessary to measure muscle performance, we measured *in vivo* the capacity of the rat to climb a 1.1 m high ladder at 80 ° inclination with increasing load with intervals of 120s in between each climb (material & methods). The experimental procedure was performed three times per week during 8 weeks. Considering the rats needed to adapt to the new environment and exercise (ladder), only trained rats were used. The data was then averaged per week and physiological and anatomical parameters were analyzed (Fig 2). The maximal capacity to carry a total load attached to the rat’s tail was measured as maximal carrying load, in which CrM supplementation significantly increased the maximal carrying load from the second week in comparison to SD-T alone. (Fig 2A, S1-Table 4). We considered the activity of climbing a ladder a similar mechanism to measuring isotonic contractions *in vitro*, with the advantage that here we were able to measure in the whole animal and not only in one single muscle. Therefore, we calculated the total isotonic force in terms of total mass (body weight plus load weight) in grams that the trained rat successfully carried to the top of the ladder. The analysis showed that, in comparison to SD-T alone, CrM supplementation significantly improved the total isotonic force from the second week of the experimental procedure and it lasted until week eight (Fig 2B, S1-Table 5). Next we evaluated whether the SD-T in combination with CrM supplementation would modify the work performed by the rats to carry up its own body weight plus an increasing load to the top of the ladder. The supplementation of SD-T rats with CrM significantly increased the work done only in the First four weeks; from the 5^th^ the to the 8^th^ week, the work was similar to SD (Fig 2C, S1-Table 6). In order to correlate the change in the muscle physiology observed in trained rats supplemented with CrM, we dissected the gastrocnemius at the end of the 8^th^ week of experiment for anatomical analyses. First, we observed that the cross-sectional area (CSA) was increased in trained rats supplemented with CrM (Fig 2D). Finally, we quantified the hypertrophic effect of CrM on trained rats measuring the distribution of the muscle fiber diameter using the minimal Feret’s diameter’ method (Fig 2E).

**Figure 2.**
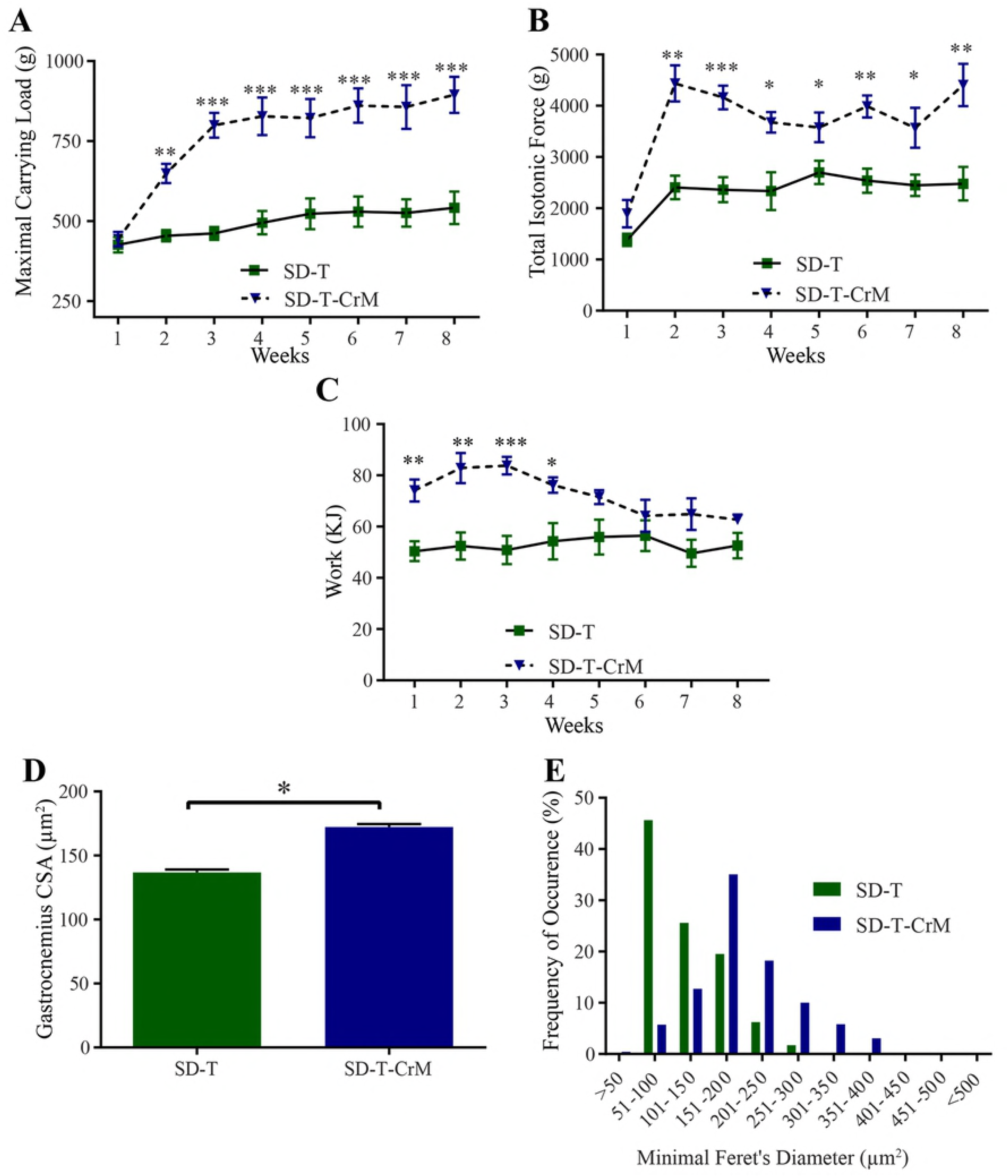
Carrying capacity, muscle performance and histological analyses on the effect of CrM supplementation (SD-T-CrM; blue-triangle) in comparison to rats without CrM supplementation (SD-T; green-square). The rats climbed the 1.1 m, 80° inclination ladder with an interval of 120 s rest in between climbing in three sessions per week during an 8 week-period. After each successful climbing attempt to the top of the ladder, the carried load was increased in 30-g steps from the starting load of 50% of the body weight. A. Maximal carrying load is the total load successfully carried to the top of the ladder. B. Effect of CrM supplementation on total isotonic contraction. Total isotonic contraction (g) was calculated by summing the body weight and the total carried load to the top of the ladder times the successful number of times the rats climbed to the top of the ladder. C. Work performance from rats receiving CrM supplementation on the climbing task over the 8 weeks of experimental procedure. D. The gastrocnemius cross-sectional area (CSA, μm^2^) from each rat was measured at the end of the 8 week-period experiment. E. Consequence of the distribution of muscle fiber diameter correction by the minimal Feret’s diameter calculation after CrM supplementation. n = 5. * p< 0.05; ** p< 0.01; *** p< 0.001.

### High-fat diet cancels the positive effect of CrM supplementation

After the characterization of the role of CrM diet supplementation on muscle performance, we evaluated whether this positive effect would be present in trained rats fed with HF diet. Under HF diet, CrM supplementation did not improve the maximal carrying load on trained rats (Fig 3A, S1-Table 7). Similarly, the total isotonic force was not improved with CrM supplemented in the diet (Fig 3B, S1-Table 8). Consequently, work performed by HF-T-CrM rats was also not different to HF-T, except on 4^th^ week, where the HF-T CrM rats had a significant increase in work done (Fig 3C, S1-Table 9). The CrM-induced hypertrophy observed in SD-T-CrM rats was absent in rats that received HF-T-CrM in comparison to HF-T rats (Fig 3D). The analysis of the muscle fiber distribution also showed no difference between HF-T and HF-T-CrM groups (Fig 3E).

**Figure 3.**
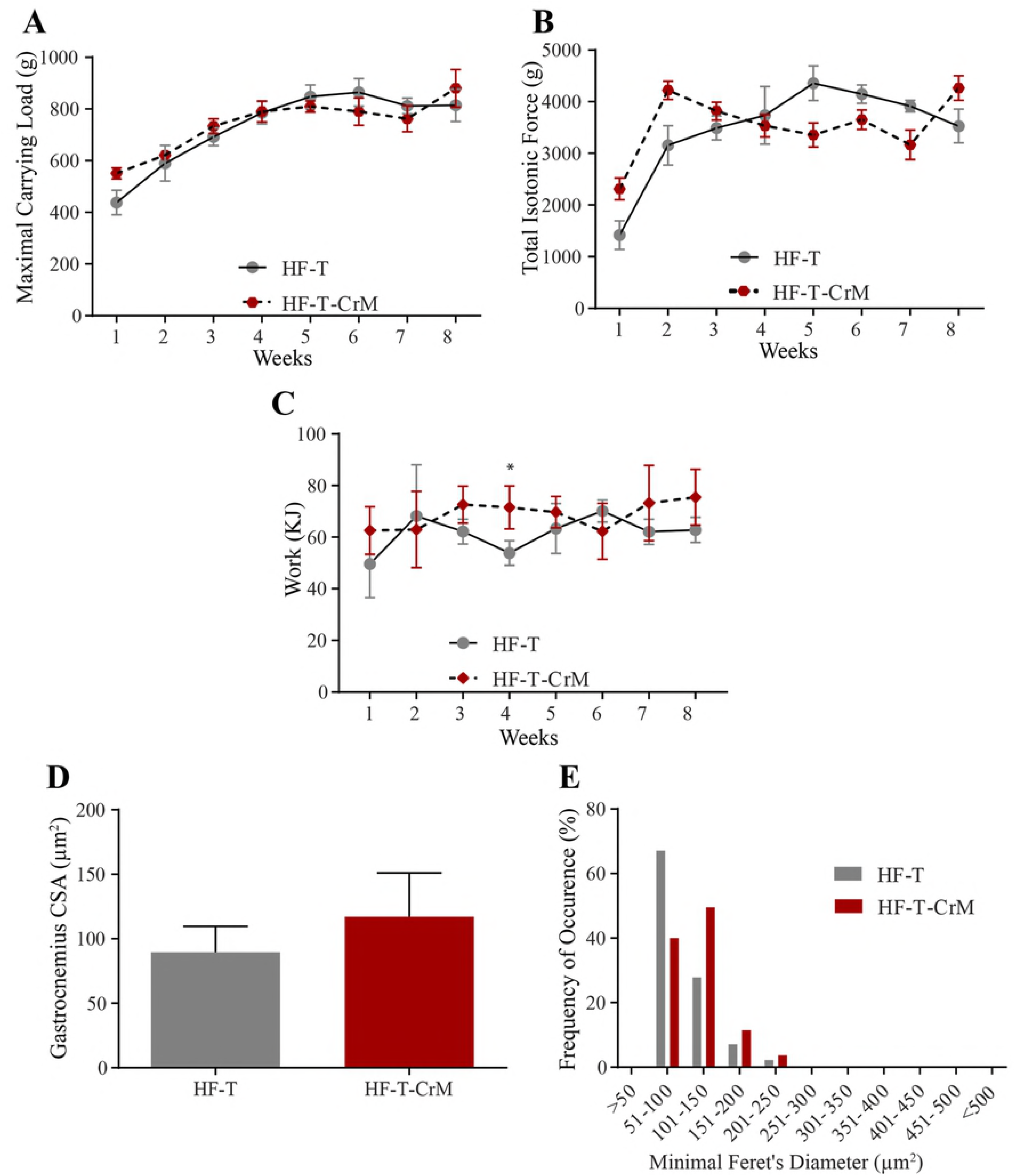
Carrying capacity, muscle performance and histological analyses on the effect of CrM supplementation (HF-T-CrM; red-hexagon) in comparison to rats under HF diet treatment (HF-T; gray-circle). The rats climbed the 1.1 m, 80° inclination ladder with an interval of 120 s rest in between climbing in three sessions per week during an 8 week-period. After each successful climbing attempt to the top of the ladder, the carried load was increased in 30-g steps from the starting load of 50% of the body weight. A. Maximal carrying load is the total load successfully carried to the top of the ladder. B. Effect of CrM supplementation on total isotonic contraction. Total isotonic contraction (g) was calculated by summing the body weight and the total carried load to the top of the ladder times the successful number of times the rats climbed to the top of the ladder. C. Work performance from rats receiving CrM supplementation on the climbing task over the 8 weeks of experimental procedure. D. The gastrocnemius cross-sectional area (CSA, μm^2^) from each rat was measured at the end of the 8 week-period experiment. E. Consequence of the distribution of muscle fiber diameter correction by the minimal Feret’s diameter calculation after CrM supplementation. n = 5. * p< 0.05; ** p< 0.01; *** p< 0.001.

### Effect of HF diet on muscle performance

Once the role of CrM supplementation was characterized within the dietary effect, we compared the outcome in muscle physiology and anatomy between SD and HF diets in trained rats. Although HF-T rats were able to carry significantly more load from the second week in comparison to SD-T rats (Fig 4A), this effect was absent upon normalization of carrying load to the body weight in the first five weeks and the effect was even significantly reversed from the 6^th^ to the 8^th^ week of the experiment (Fig 4B). The total isotonic force was significant higher in HF-T rats in comparison to SD-T rats (Fig 4C); however, this difference was absent once the total isotonic force was normalized by the body weight (Fig 4D). The analyses of work (Fig 4E) and work normalized by body weight (Fig 4F) showed that there is no difference between SD-T and HF-T groups.

**Figure 4.**
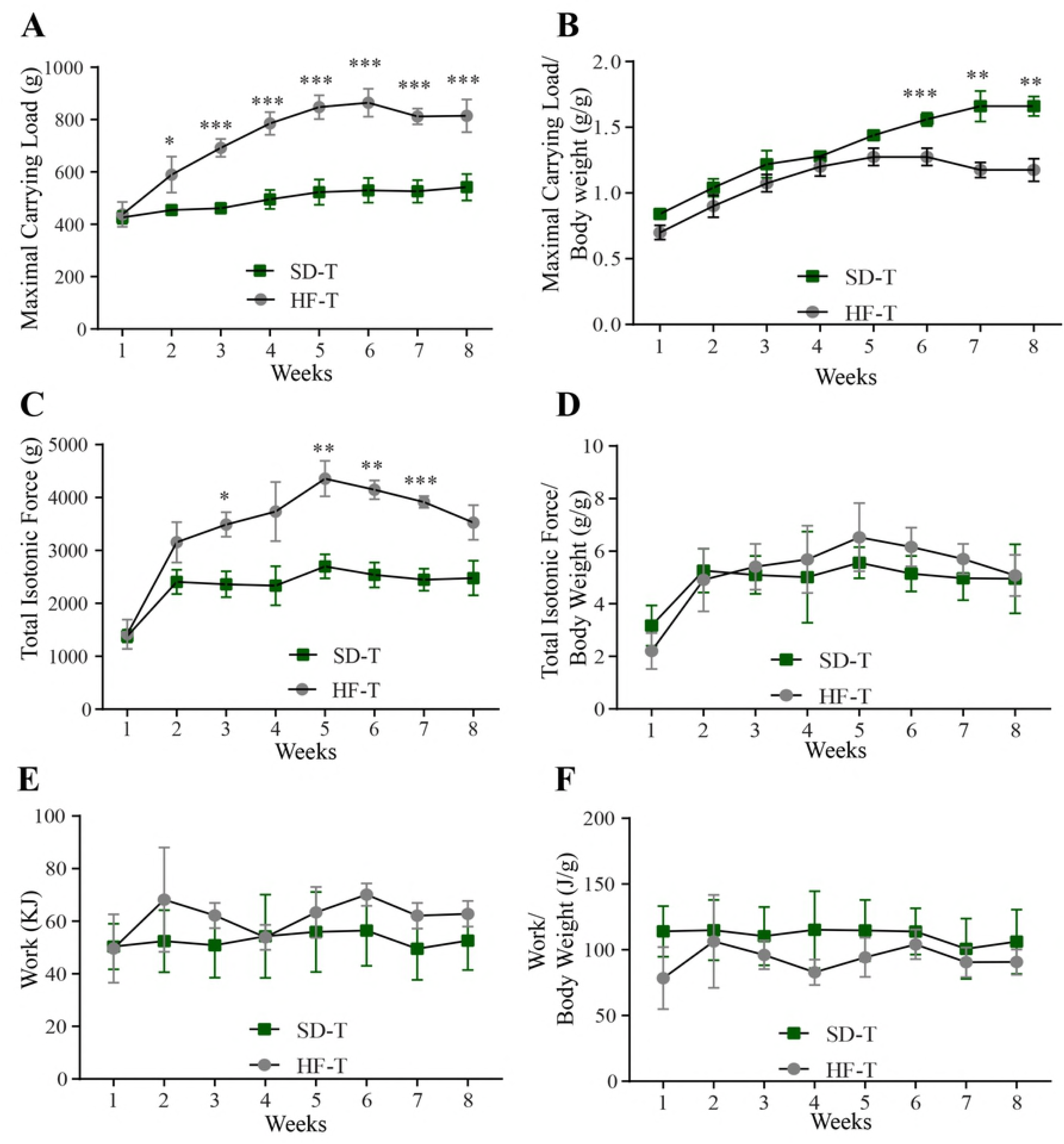
Comparison of the diet’s effect on muscle performance between SD-T rats (green-square) and HF-T rats (grey-circle). A. Maximal carrying load (g). B. Carrying load normalized to body weight (g/g). C. Total isotonic contraction (g). D. Total isotonic contraction normalized to body weight (g/g). E. Work (KJ). F, Work normalized to body weight (KJ/g). n = 5. * p< 0.05; ** p< 0.01; *** p< 0.001.

### Positive role of CrM supplementation is diet-dependent

CrM supplementation is already known to have ergogenic effects on skeletal muscles; therefore, we investigated whether CrM would improve the muscle performance in HF- rats in comparison to SD-T CrM rats. Although there was no difference in maximal carrying load between SD-T-CrM and HF-T-CrM rats (Fig 5A), we observed that from the 2^nd^ until the 8^th^ week of training, muscle performance in SD-T-CrM rats was significantly improved once the maximal carrying load was normalized by the body weight (Fig 5B). Although the analyses of total isotonic force (Fig 5C), total isotonic force normalized by the body weight (Fig 5D), work (Fig 5E) and work normalized by the body weight (Fig 5F) showed that SD-T CrM rats had better performance, but this was not significant in comparison to HF-T-CrM rats.

**Figure 5.**
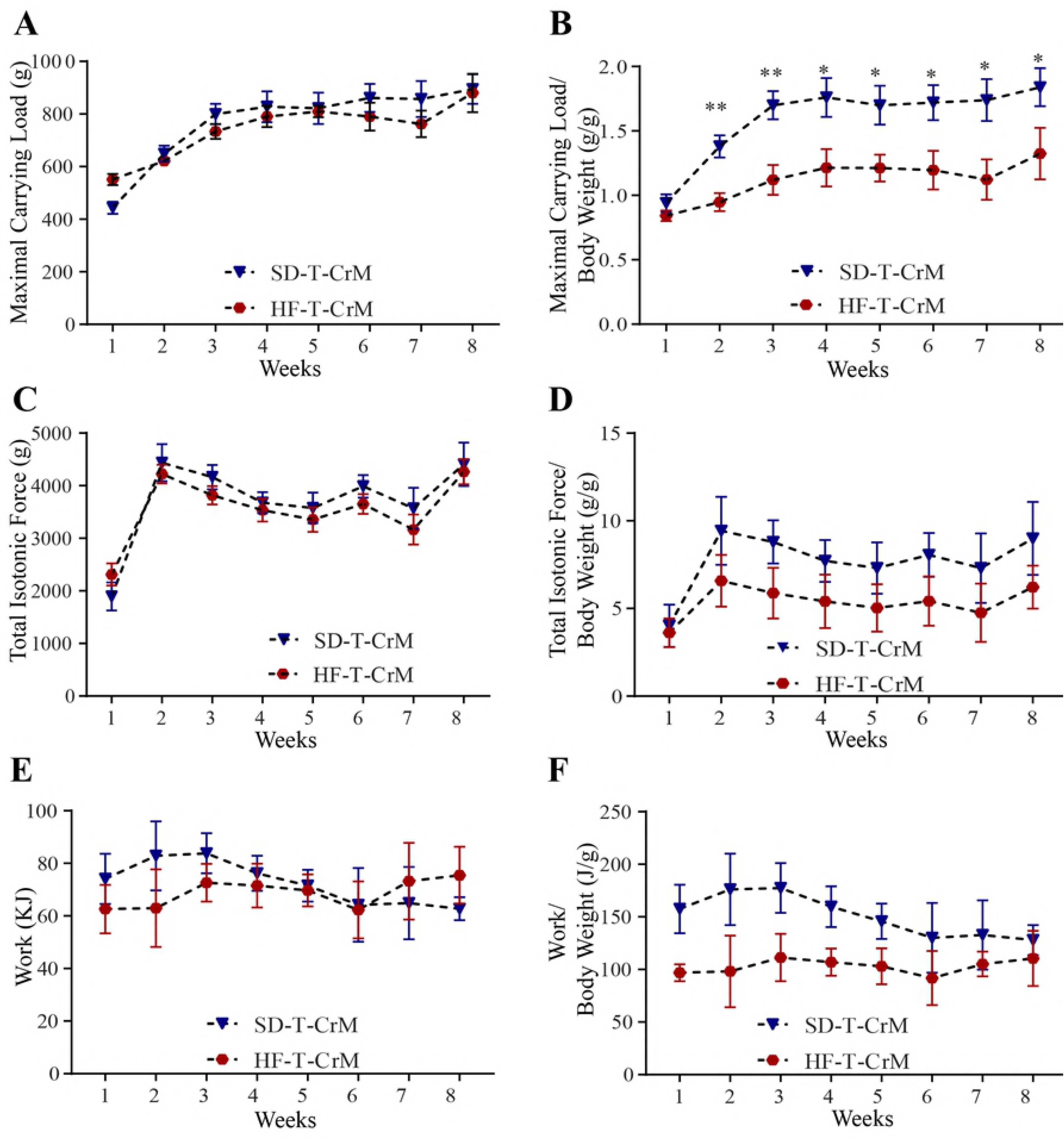
Comparison of the CrM supplementation effect on muscle performance between SD-T rats (blue-triangle) and HF-T rats (red-circle). A. Maximal carrying load (g). B. Carrying load normalized to body weight (g/g). C. Total isotonic contraction (g). D. Total isotonic contraction normalized to body weight (g/g). E. Work (KJ). F, Work normalized to body weight (KJ/g). n = 5. * p< 0.05; ** p< 0.01.

### Dysregulation of the IGF-PI3K-AKT-mTOR signaling pathway in HF-T rats

The data presented above suggested that CrM improved muscle performance, CSA and distribution of fiber diameter in trained rats fed with SD, but this effect was absent in trained rats fed with HF diet. Thus, we hypothesized that the main muscle synthesis pathway, the IGF-PI3K-AKT-mTOR signaling pathway would be enhanced in SD-T-CrM rats in comparison to SD-T rats and this effect would be reduced in rats fed with HF diet. Using gastrocnemius muscle samples, we quantified the normalized protein level of IGF, observing that trained rats fed with SD and supplemented with CrM significantly increased IGF level in comparison to SD-T rats, but HF significantly reduced IGF levels in both HF-T and HF-T-CrM rats (Fig 6A). Next, we quantified the protein level of β IGF receptor subunits, and as observed for IGF, SD-T-CrM rats showed a significant increase in comparison to SD-T rats (Fig 6B); Again, HF diet reduced protein levels of β IGF receptor subunits in comparison to SD-T rats. The HF-T-CrM also showed a non-significant reduction in IGFR levels in comparion to SD-T-CrM (Fig 6B). As the insulin receptor substrate 1 (IRS1) is the next step of the muscle synthesis pathway, the protein level analysis showed that subunit β did not change in rats receiving SD, but in comparison to SD-T rats, it was significantly reduced in rats fed with HF diet (Fig 6C). We next evaluated whether CrM would modify protein levels of PI3K; although CrM did not change the protein levels under SD-T treatment, HF-T significantly reduced the PI3K protein levels in comparison to SD-T. We analyzed protein levels of AKT, one of the key elements of the pathway, as total AKT, phosphorylated AKT and the ratio of total and phosphorylated AKT. Although the normalized to GAPDH protein levels of total and phosphorylated AKT were not different among the groups (data not shown), there was a significant difference between the ratio between the phosphorylated AKT normalized to GAPDH (phospho-AKT/GAPDH) and the total AKT normalized to GAPDH (Total AKT/GPADH). While phospho-AKT/Total AKT was significantly higher in SD-T-CrM in comparison to SD-T rats, it was significantly lower in HF-T-CrM rats in comparison to SD-T-CrM rats (Fig 6E). Analyzing the effect of diet and CrM at the mTOR protein level, we observed that there was no difference in total mTOR normalized to GAPDH among the treatments (Fig 6F); however, HF-T treatment significantly reduced the normalized protein levels of phosphorylated mTOR in comparison to SD-T and in HF-T-CrM in comparison to SD-T-CrM (Fig 6G). This tendency was also observed in the ratio between phosphorylated mTOR and total mTOR (Fig 6H). Finally, we analyzed the role of HF diet at the protein levels of S6K, one of the final effectors of the pathway, showing that only HF-T-CrM significantly reduced the protein levels in comparison to SD-T-CrM (Fig 6I). The above results suggested that CrM supplementation in standard diet might overexpress the ergogenic IGF1 pathway, but HF diet significantly reduced the beneficial effect of CrM.

**Figure 6.**
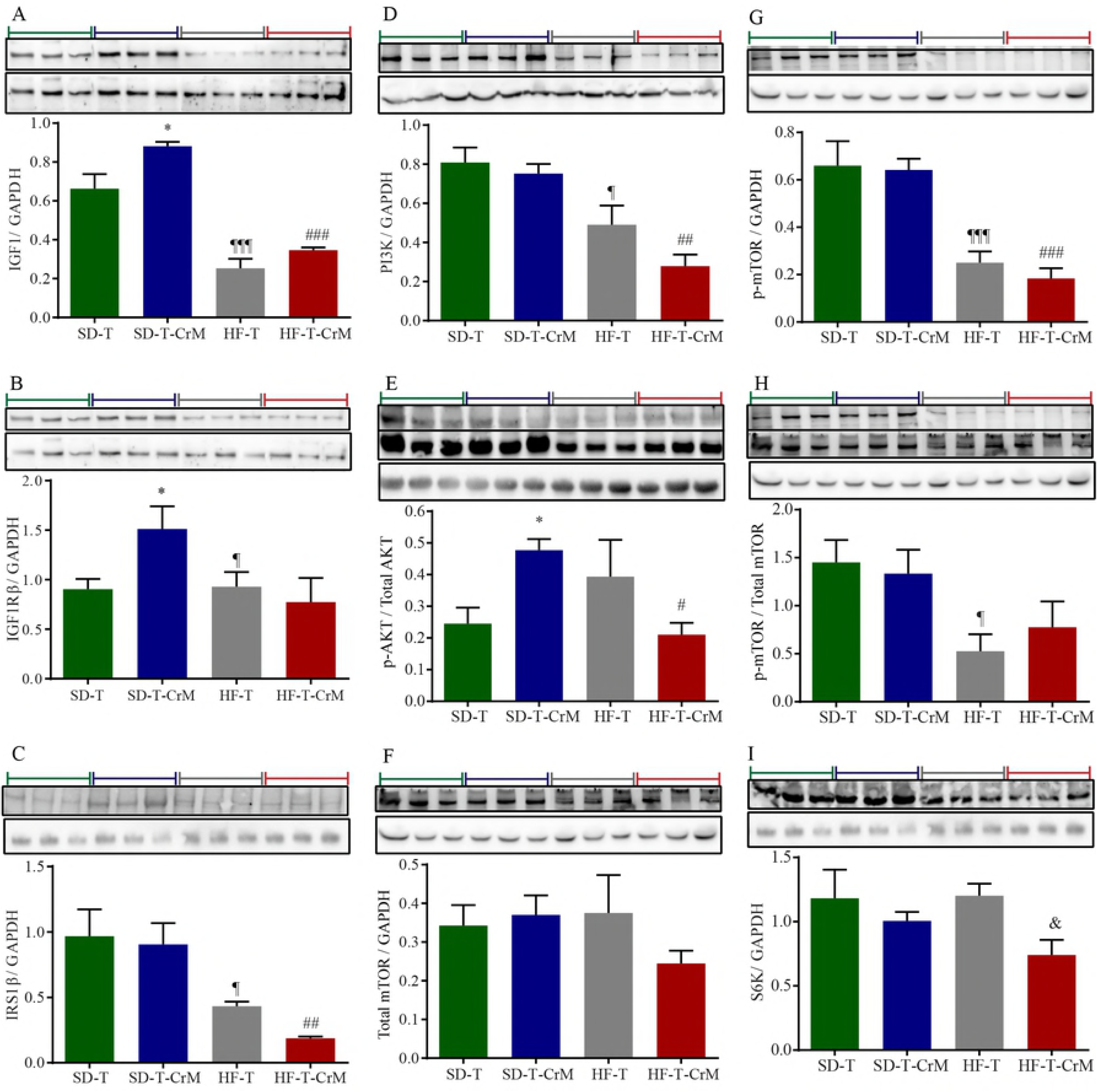
Immunoblotting analyzes of protein expression of IGF-1-PI3K-AKT-mTOR pathway from gastrocnemius muscle. The protein levels of GAPDH shown at the bottom of each immunoblot were used to normalize by the protein levels of each immunoblot shown at the top. Three independent experiments are shown and separated according to the treatment, indicated with a color-coded bar; green represents SD-T group, blue represents SD-T-CrM group, grey represents HF-T group and red represents HF-T-CrM group. A. IGF-1. B. IGF-1 receptor β subunit. C. IRS1 β subunit. D. PI3K. E. Phosphorylated AKT. F. Total mTOR. G. Phosphorylated mTOR. H. Ratio phosphorylated mTOR to total mTOR. I. S6K. n = 3. * p< 0.05 *vs* SD-T; ¶ p< 0.05 *vs.* SD-T; ¶¶¶ ** p< 0.001 *vs*. SD-T; # p< 0.05 *vs*. SD-T-CrM; ## p< 0.01 *vs*. SD-T-CrM; ### p< 0.001 *vs*. SD-T-CrM; & p< 0.05 *vs*. HF-T-CrM.

## Discussion

Using resistance ladder-climbing training as a model to measure muscle performance, we showed that HF diet impairs muscle performance by inhibiting protein expression of the IGF1-IRS1-PI3K-AKT-mTOR pathway. In comparison to SD this effect was not rescued with the supplementation of CrM in the diet. We observed that HF diet was the main factor of increased body weight, mainly due to a significant increase of epididymal fat mass, instead of other possible factors, such as lack of exercise or CrM supplementation only. Therefore, we focused our study on the effect of diet (SD *vs* HF diet) and evaluated how CrM would improve the muscle performance under different diets. Instead of using a high number of animals for the analyses of muscle performance *in vitro*, we used the strategy to train the rats to the resistance ladder-climbing training model (1.1 m, 80° of inclination, 120s of rest in between climbing) carrying progressively increasing loads (30g increases starting from 50% of body weight). The progress of the treatment in each rat was measured and analyzed over 8 weeks, thus reducing the number of animals required for the statistical analyses. The parameters analyzed were maximal carrying load, total isotonic force and work, providing a complete outcome of CrM supplementation on the diet over the rat’s general fitness. During the eight weeks of resistance ladder-climbing training, we observed that CrM supplementation to SD significantly increased the rat’s capacity to climb the ladder with increasing load (Fig 2). This increase was associated with muscular hypertrophy (Fig 2D and 2E). This result was supported by protein level analyses showing that CrM supplementation on SD increased IGF and phosphorylated AKT (Fig 6). Conversely, under HF diet CrM supplementation was not able to improve muscle performance measured as maximal carrying load, total isotonic force and work under HF diet, without the expected muscle hypertrophic effect (Fig 3) due to inhibition of protein expression of IGF1-IRS1-PI3K-AKT-mTOR pathway. All together, these results suggested that HF is the major negative effect on muscle performance measured by the resistance ladder-climbing training model attributable to inhibition of exercise- and CrM supplementation- mediated muscle protein synthesis.

### CrM supplementation improves muscle performance under SD

Creatine monohydrate (CrM) supplementation is the most common nutritional supplement used by athletes in combination with resistance exercise and more recently in elderly patients to avoid sarcopenia [20, 21]. Creatine can be obtained from meat, but also can be synthetized in the body from arginine, glycine and methionine and its main function as phosphocreatine, is to buffer adenosine triphosphate levels in the muscle improving and enhancing muscle performance during exercise [22, 23]. In a recent meta-analysis, it has been shown that CrM supplementation during resistance training increased lean tissue mass by ca. 1.4kg resulting in a significant increase in force in comparison to placebo [24]. The possible mechanism that creatine increases muscle mass and force is increasing the expression of insulin-like growth factor-1 (IGF-1) [16, 25], which would activate the key elements of protein synthesis of the IGF1-IRS1-PI3K-AKT-mTOR pathway [17, 26, 27]. The resultant increase of IGF-1 via creatine is also observable in the significantly increased expression of several myogenic regulatory factors, such as Myo-D, Myf-5 and MRF-4 (Luois 2004), which are responsible for synchronized triggering of satellite cell activation, proliferation and differentiation (Zanou 2013). This positive effect of creatine on muscle is probably only observable together with exercise [28–30].

We observed that rats under standard diet in combination with training and CrM supplementation significantly increased maximal carrying load, total isotonic force and work in comparison to the SD-T rat group. We also observed that the SD-T-CrM group had significantly higher gastrocnemius CSA, with a shift to the right on the minimal Feret’s diameter (Fig 2). These results support previous reports showing that CrM is responsible to improve the effects of resistance training on muscle performance via the IGF1-IRS1-PI3K-AKT-mTOR pathway.

### The positive effect of resistance training and CrM are inhibited by HF diet

It has been extensively shown in humans and animal models that long term HF diet results in an excessive accumulation of adipose tissue in skeletal muscle leading to muscle atrophy via activation of proteins of the atrophy pathway (TNFα-TNF-R-NFκB-MuRF-1); as consequence, not only body weight increases but also the ubiquitin proteasome system, autophagy, and apoptosis pathways are activated [2–4, 31–34]. Another reported consequence of HF diet, is a reduction in muscle diameter, specific force and thus percentage of muscle strength [4]. It has also been described that long term HF diet impairs all described benefits of resistance training by reducing cortical actin filaments, impairing insulin stimulated glucose transport, reducing matrix metallopoteinases activity and reducing IRS-1 Pi3K kinase activity [35–38]. The main strategy to recover muscle force after HF diet is through resistance training which causes increased expression of contractile proteins and the muscle glucose transporter 4. Further, resistance training increases IRS-1 Pi3K kinase activity resulting in activation of AKT kinase, thus improving muscle performance [36, 37, 39]. Our results support and extend previous findings showing that HF diet significantly increased body weight and epididymal fat mass. In comparison to SD, our data from the HF group initially suggested an increase of maximal carrying load and total isotonic force; however, once these parameters were normalized to the body weight, rats from the SD group were able to carry significantly more load than those from the HF group, with no difference in total isotonic force (Fig 4). These results are supported by the inverse relationship between fiber size and loss in force generation capacity in *in vitro* muscle fibers in obese older mice [40, 41]. Moreover, our protein level expression analyses suggested that HF diet significantly reduced main targets of the protein synthesis in almost the entire IGF1-IRS1-PI3K-AKT-mTOR pathway (Fig 6).

Differently from what we expected, CrM supplementation was not able to overcome the negative effect of HF diet on the parameters evaluated here in terms of muscle performance (Fig 3). Moreover, by comparing SD-T-CrM and HF-T-CrM data normalized to body weight (Fig 5), we observed that diet has a crucial effect on muscle performance. Long-term HF diet changes muscle fiber-type and myofibers inhibiting thus the CrM and resistance training effects on muscle contractile force [41]. Once we evaluated the protein expression, we observed a similar protein expression of HF-T-CrM in comparison to HF-T group, with the interesting difference that the HF-CrM group showed significantly reduced protein levels of phospho-AKT and S6K. The detailed analysis of the protein levels shown in Fig 6 and summarized in Fig 7, suggested specific action of CrM, HF and HF-CrM in the IGF 1-IRS 1-PI3K-AKT-mTOR pathway of trained rats. Interestingly, largely in accordance with previous reports, CrM did not activate all elements of the IGF1-IRS1-PI3K-AKT-mTOR pathway, but rather mainly the expression of IGF-1 and, the novelty here, additionally the expression of phospho-AKT. Conversely, HF diet inhibits the expression of several proteins of the IGF1-IRS1-PI3K-AKT-mTOR pathway, such as IGF-1 receptor, IRS1, PI3K, mTOR, as well as IGF-1, which in turn was activated by CrM. Impressively, the supplementation of CrM on HF diet did not revert the HF diet-induced down-regulation, but also included down-regulation of phospho-AKT and S6K, another two major elements of the IGF1-IRS1-PI3K-AKT-mTOR pathway (Fig 7).

**Figure 7.**
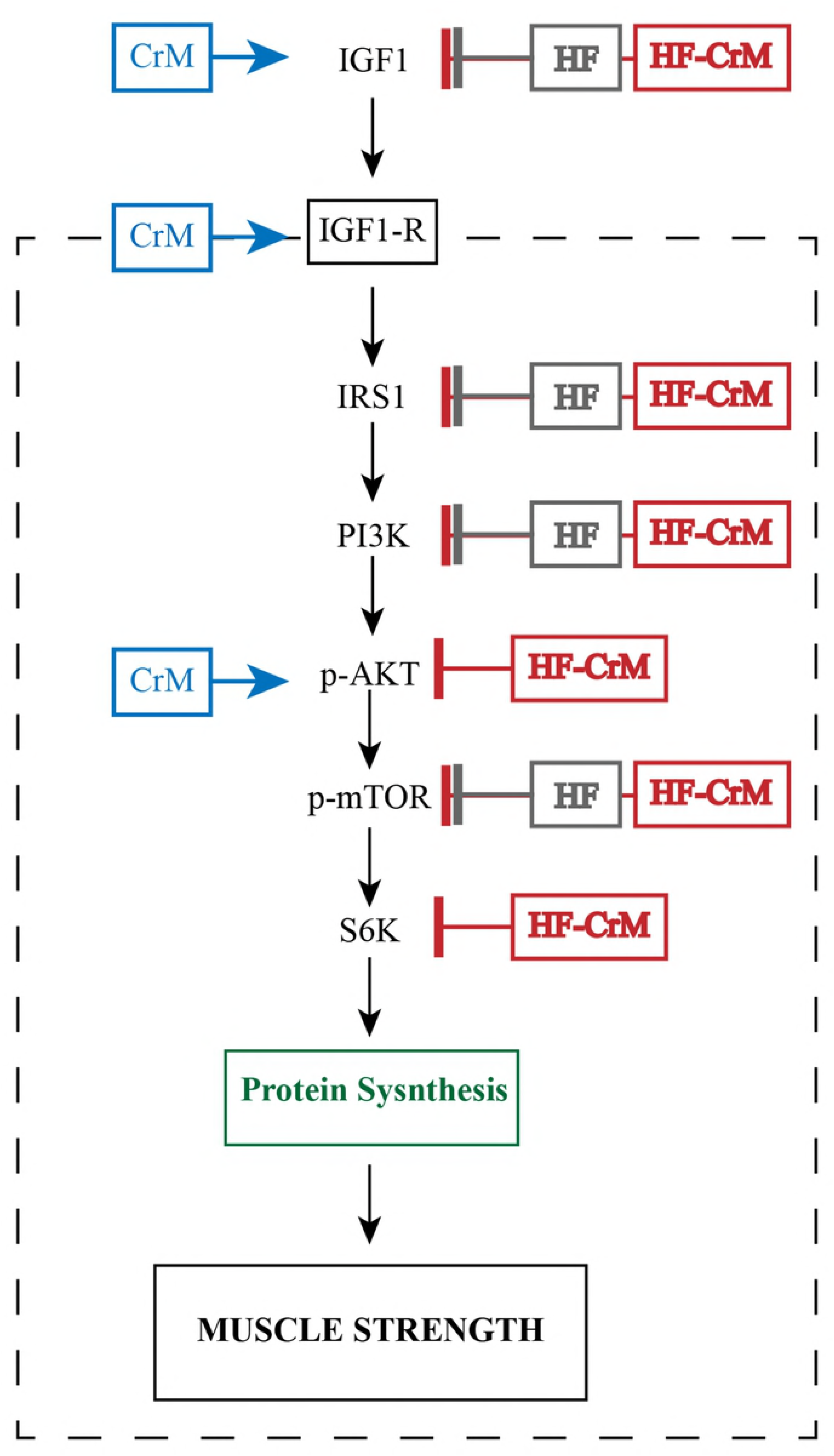
Summary of the effect on the protein levels of the gastrocnemius muscle of the IGF-1-PI3K-AKT-mTOR pathway after 8 weeks of resistance training and receiving CrM, HF or HF-CrM. Creatine monohydrate supplementation (CrM) increased the protein levels of IGF1, IGF1-receptor and phosphorylated AKT. This enhancement of the protein levels promoted by CrM supplementation would explain the increase of muscle performance. High-fat diet (HF) and high-fat diet supplemented with CrM (HF-CrM) decreased the protein levels of IGF1, IRS1, PI3K, phosphorylated mTOR, and HF-CrM also decreased the protein level of phosphorylated SKT and S6K. The decrease on the protein level of these key targets of IGF-1-PI3K-AKT-mTOR pathway would explain the reduced muscle performance.

## Conclusion

We demonstrated the mechanism by which during resistance training CrM increases muscle size and muscle performance, suggesting a higher activation of muscle protein synthesis via IGF1-IRS1-PI3K-AKT-mTOR pathway. Conversely, HF diet reduces muscle size and performance by inhibiting the expression of the same IGF1-IRS1-PI3K-AKT-mTOR pathway and this effect was not overruled by supplementation of CrM. These results suggested the necessity to change the diet prior in order to perceive the benefit of resistance training and CrM supplementation.

## Acknowledgements

Luiz Gustavo de Almeida Chuffa and Luis Antonio Justulin Junior for kindly providing the antibodies. Bruno Ricardo Santos da Silva for his technical assistance resistance training protocol and preparation of tissue for histology. Carlos Alberto da Silva for the access to the ladder-climbing equipment. Ulrike Zeiger for her valuable comments in the manuscript.

## Support information

**S1 Table 1. List of antibodies used on immunoblotting.**

**S1 Table 2. Comparison of the effect of standard diet (SD) and high-fat diet (HF) on body end of the 8^th^ weight (g) at the week of experiment.**

**S1 Table 3. Comparison of the effect of standard diet (SD) and high-fat diet (HF) on epididymal fat mass (g) at the end of the 8^th^ week of experiment.**

**S1 Table 4. Summary of the statistical analysis for maximal carrying load (g) between SD-T and SD-T-CrM.** The maximal carrying load was calculated from the total amount of load carried to the top of the ladder.

**S1 Table 5. Summary of the statistical analysis for total isotonic force (g) between SD-T and SD-T-CrM.** The value was calculating the sum of body weight plus maximal carrying load times the successful times the animal climbed the ladder.

**S1 Table 6. Summary of the statistical analysis for work (kJ) between SD-T and SD-T-CrM.** Work was calculated multiplying total mass lifted to the top of the ladder, the length of the ladder (1.1m), gravitational force (9.8 06 ms^−2^) and the ladder’s angle (sen80 = 0.9848).

**S1 Table 7. Summary of the statistical analysis for maximal carrying load (g) between HF-T and HF-T-CrM.** The maximal carrying load was calculated from the total amount of load carried to the top of the ladder.

**S1 Table 8. Summary of the statistical analysis for total isotonic force (g) between HF-T and HF-T-CrM.** The value was calculating the sum of body weight plus maximal carrying load times the successful times the animal climbed the ladder.

**S1 Table 9. Summary of the statistical analysis for work (kJ) between HF-T and HF-T-CrM.** Work was calculated multiplying total mass lifted to the top of the ladder, the length of the ladder (1.1m), gravitational force (9.8 06 ms^−2^) and the ladder’s angle (sen80 = 0.9848).

